# Averaging a local PLSR pipeline to predict chemical compositions and nutritive values of forages and feed from spectral near infrared data

**DOI:** 10.1101/2023.08.16.553515

**Authors:** Matthieu Lesnoff

**Affiliations:** SELMET, Univ Montpellier, CIRAD, INRAe, Institut Agro, Montpellier, France; CIRAD, UMR SELMET, Montpellier, France; ChemHouse Research Group, Montpellier, France

**Author notes:** Corresponding author: Selmet Joint Research Unit (Tropical and Mediterranean Animal Production Systems), Cirad, TA C-112 / A - Campus international de Baillarguet - 34398 Montpellier Cedex 5, France.

**Keywords:** local PLSR, *k*-nearest neighbors, model averaging, NIR data, forages

## Abstract

Partial least squares regression (PLSR) is a reference method in chemometrics. In agronomy, it is used for instance to predict components of chemical composition (response variables ***y***) of vegetal materials from spectral near infrared (NIR) data ***X*** collected from spectrometers. The principle of PLSR is to reduce the dimension of the spectral data ***X*** by computing vectors that are then used as latent variables (LVs) in a multiple linear model. A difficulty is to determine the relevant dimensionality (number of LVs) of the model for the given available data. This step can also become time consuming when many different datasets have to be processed and/or the datasets are frequently updated. An alternative to determinate the relevant PLSR dimensionality is the ensemble learning method “PLSR averaging”. In the past, this method has been demonstrated to be efficient for complex biological materials such as mixed forages, and facilitates to automatize predictions (e.g. in user-friendly web interface platforms). This article presents the extension of the PLSR averaging to a *k*-nearest neighbors locally weighted PLSR pipeline (kNN-LWPLSR). The kNN-LWPLSR pipeline has the advantage to account for non-linearity between **X** and **y** existing for instance in heterogeneous data (e.g. mixing of vegetal species, collection from different geographical areas, etc.). In the article, kNN-LWPLSR averaging is applied to an extensive NIR database built to predict the chemical composition of European and tropical forages and feed. The main finding of the study was the overall superiority of the averaging compared to the usual kNN-LWPLSR. Averaging may therefore be recommended in local PLSR pipelines to predict NIR forage and feed data.

## 1 Introduction

Near-infrared spectroscopy (NIRS) is a fast and nondestructive analytical method used in many agronomic contexts, for instance to evaluate the nutritive quality of forages. Basically, spectral data ***X*** (matrix of *n* observations × *p* wavelengths) are collected on samples of the material to study (e.g. forages) using a spectrometer, and targeted response variables (e.g. chemical compositions) ***Y*** {***y***_1_, …, ***y***_*k*_} (*k* vectors of *n* observations) are measured precisely in laboratory. Regression models of ***Y*** on ***X*** are then fitted and used to predict the response variables from new spectral observations. Spectral data are known to be highly collinear and, in general, matrix ***X*** is ill-conditioned. Specific regression methods have to be implemented, in particular partial least squares regression (PLSR) [1– 3]. The general principle of PLSR is to reduce the dimension of ***X*** to a limited number *a* << *p* of orthogonal vectors *n* × 1 maximizing the squared covariance with ***Y*** and referred to as scores. The scores are then used as regressor latent variables (LVs) in a multiple linear regression (MLR). PLSR is very efficient when the relationship between ***X*** and ***Y*** is linear [4].

For several years, agronomic databases (e.g. in feed, food or soils researches) tend to aggregate large numbers of samples of different natures or origins, bringing heterogeneity. This generates curvatures and/or clustering in the data that can alter the linear relation between ***X*** and ***Y*** and therefore the PLSR predictions. Local PLSR is an easy tool that can turn out non-linearity in the data [4–7]. The general principle is, for each new observation to predict, to do a pre-selection of *k* nearest neighbors of the observation (the kNN selection step) and then to apply a PLSR to the neighborhood (i.e. the *k* neighbors). Many variants of local PLSR pipelines can be built, depending essentially on the type of PLSR implemented, and on how are selected the neighborhood. One of these variants ([8]) consists in applying a locally weighted PLSR (LWPLSR) on the neighborhood, instead of a PLSR as it is done in the more common local PLSR, say kNN-PLSR. LWPLSR, detailed in a next section, has the particularity to weight each of the *n* training observations {***x***_*i*_; *i* = 1,…,*n*} depending on its distance (or any dissimilarity) to the observation to predict, ***x***_new_ (while in PLSR a uniform weight 1 / *n* is given to all the ***x***_*i*_). Closer is ***x***_*i*_ to ***x***_new_, higher is its weight in the iterative PLSR equations and therefore its importance in the prediction. Implementing LWPLSR in the local pipeline, say kNN-LWPLSR ([8]), has been observed to be more efficient than using kNN-PLSR for various data including forages, for regression as for discrimination [8,9]. As a remark, kNN-LWPLSR can be considered as a particular case of LWPLSR where positive weights are given to the neighbors of ***x***_new_ and null weights to the observations outside of the neighborhood. Nevertheless, for large datasets, doing the kNN step and then apply LWPLSR only on the neighborhood (kNN-LWPLSR) is much faster in terms of computation times than implement LWPLSR on the all data ([8]).

An important step of tuning kNN-LWPLSR, as for PLSR and LWPLSR, is to determine the dimensionality (i.e. the number of LVs), say *a*, that is used in the MLR step. Different strategies have been addressed in the PLSR literature to guide the determination of an optimal dimensionality [10–15]. One of the most common is the cross-validation (CV) that searches the value *a* that minimizes the CV-error curve. These strategies face to several difficulties. Firstly, despite attempts of automated procedures, they often require case-by-case decisions based on expertise. Secondly, to find a unique optimal dimensionality is in general difficult when the data contain complex information, which is often the case with biological materials such as plants and forages (mixing of stems, leaves, different stages of development and geographical areas, etc.). Finally, finding *a* can also become time consuming when many datasets have to be processed (e.g. many variable responses ***y*** to consider successively) and/or when the datasets are periodically updated with new training observations. Alternatively, PLSR-averaging [16–19] (say PLSR-AVG) is an ensemble learning method whose the main objective is to bypass the determination of *a*. The method consists in averaging the predictions of *A* + 1 PLSR models of dimensionality *r* = 0, 1, 2 … *A* LVs. The maximal value *A* is set *a priori* and voluntary larger than the expected values of optimal *a*. For forages ([19]), PLSR-AVG was shown more efficient than the usual procedure where *a* was determined by CV. In the past, such an averaging has already been implemented in a kNN-PLSR pipeline ([18]). The present article proposes its extension to the kNN-LWPLSR pipeline [8].

The article is organized as follows. Theoretical points on kNN-LWPLSR and PLSR-AVG are firstly presented. Then, the kNN-LWPLSR-averaging pipeline (say kNN-LWPLSR-AVG) is applied to an extensive NIR database on chemical composition of European and tropical forages and feed. The predictive performance of the pipeline is compared to kNN-LWPLSR where the optimal number of LVs *a* is determined by CV.

## 2 Theory

### 2.1 kNN-LWPLSR

LWPLSR and kNN-LWPLSR have been described in Lesnoff et al. [8]. LWPLSR [20– 22] is a particular case of weighted PLSR (WPLSR). In WPLSR, a *n* × 1 vector of weights *δ* = {*δ*_1_, *δ*_2_, … *δ*_*n*_} is inserted into the PLSR algorithm, in two steps: (a) the PLS scores (LVs) are computed by maximizing weighted (instead of unweighted) squared covariances, and (b) the prediction MLR equation is computed by weighted least-squares (WLS, instead of ordinary LS) on the scores. The specificity of LWPLSR compared to WPLSR is that *δ* is computed from a decreasing function, say *f*, of the distances (or any dissimilarities) between the *n* training observations and ***x***_new_. This is the same principle as in the well-known locally weighted regression algorithm [23,24]. kNN-LWPLSR simply adds a preliminary step to LWPLSR: a neighborhood is selected around ***x***_new_ (kNN selection step) on which LWPLSR is then applied.

This present article uses the same kNN-LWPLSR pipeline as in Lesnoff et al. [8], involving a fast PLSR algorithm [25] and consisting in the following steps. Firstly, a global PLSR (i.e. over all the training observation) is fitted and defines a global score space. Secondly, for each new observation ***x***_new_, Mahalanobis distances in this global score space between the training observations and ***x***_new_ are computed (obviously, other global spaces, e.g. PCA or nonlinear kernel-PCA/PLS score spaces, and/or other types of distances or dissimilarities could be used). Theses distances are used to compute the neighborhood of ***x***_new_ (kNN selection) and the weights *δ* within the neighborhood. The weight function, *f*, chosen to be easily tunable, has a negative exponential shape whose the sharpness depends on a scalar parameter, say *h* [8,21]: lower is *h*, sharper is function *f* and therefore more the closest neighbors of ***x***_new_ have importance in the LWPLSR fit. The case *h* = ∞ is the unweighted situation corresponding to a kNN-PLSR.

### 2.2 PLSR-averaging

Let ***x***_new_ be a new observation to predict, and y_new,A_ its PLSR prediction when the dimensionality (nb. LVs) is *A*. The PLSR-AVG prediction is defined by:

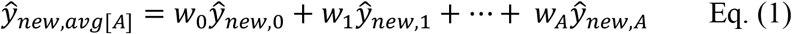

where *w*_*r*_ (*r* = 0, …, *A*) is the weight (bounded between 0 and 1) of the model with *r* LVs, with the constraint:

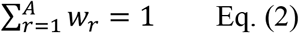

In Eq. (1), the particular case y_new,O_ is the simple mean of ***y***. Vector ***w*** = {*w*_0_, *w*_1_, …, *w*_*A*_} represents the pattern of weights, and the shape of this pattern is specific to a given averaging method. In practice, weight *w*_*r*_ should quantify the level of confidence that can be awarded to the PLSR model with *r* LVs, relatively to the other dimensionalities. Vector ***w*** can be for instance estimated from the predictive performance of each of the *A* + 1 PLSR models (e.g. estimated by CV, Akaike criterion or other indicators; [19]) or from more integrated Bayesian approaches ([26,27]).

A close approach to model averaging is the stacking ([26]) in which weights *w*_*r*_ (*r* = 0, …, A) are estimated from a meta regression model, are not bounded to [0, 1] and constrained to sum to 1. The stacking did not show improvements in the prediction performances of forages composition ([19]) compared to averaging and is not considered in this study.

### 2.3 kNN-LWPLSR-AVG

The proposed kNN-LWPLSR-AVG pipeline consists simply in chaining kNN-LWPLSR and the PLSR-AVG principle: for each neighborhood, the predictions returned from the LWPLSR models with dimensionalities *r* = 0, …, *A* are averaged.

Preliminary results on forages ([19]) showed that the uniform weighting (i.e. *w*_*r*_ = 1 / (A + 1); *r* = 0, …, *A*) was in general as efficient as more elaborated patterns on such data. Therefore, for simplicity, the present article only considers the uniform weighting that has also the advantage to be much faster to compute than other patterns. Speed is essential when implementing local PLSR pipelines since one model per new observation to predict (***x***_new_) has to be fitted.

## 3. Material and Methods

### 3.1 Dataset

The predictive models were assessed on a NIR database built by Cirad (Mediterranean and tropical livestock systems research unit) on European and tropical forages and feed. The absorbance spectra were collected on dried and grounded materials using Foss Instruments 5000 or 6500 models in the spectral range 1100 to 2498 nm (2 nm steps). The objective of the database is to enable the prediction of twelve main variables of chemical composition (Table 1). The data are regularly updated with new observations collected from various Cirad research projects and partners. The total size of the dataset used in this study was *N* = 18813 observations but, depending on the availability of the response variable, data size ranged from *N* = 2020 observations to *N* = 18055 observations (Table 1).

**Table 1:** Variables of forage and feed chemical composition to be predicted (*N* = total number of observations).

| Abbreviation | Unit | Response variable | $N$ | Mean | Min | Max | Std |
| --- | --- | --- | --- | --- | --- | --- | --- |
| DM | % | Dry matter | 18055 | 92.2 | 61.4 | 99.9 | 2.5 |
| ASH | %DM | Mineral matter | 17639 | 9.7 | 0.04 | 99.9 | 8.3 |
| CP | %DM | Crude protein | 16640 | 14.9 | 0.20 | 98.4 | 12.3 |
| EE | %DM | Crude fat | 7143 | 5.7 | 0.08 | 69.4 | 7.2 |
| CF | %DM | Crude fiber | 13232 | 25.9 | 0.06 | 77.7 | 11.9 |
| NDF | %DM | Neutral detergent fiber | 10677 | 52.0 | 0.60 | 95.1 | 18.1 |
| ADF | %DM | Acid detergent fiber | 11360 | 33.3 | 0.14 | 89.8 | 13.5 |
| ADL | %DM | Acid detergent lignin | 11072 | 8.3 | 0.01 | 50.8 | 6.7 |
| DMDCELL | %DM | DM enzymatic digestibility | 9187 | 52.5 | 6.98 | 99.9 | 17.9 |
| OMDCELL | %DM | OM enzymatic digestibility | 8857 | 50.4 | 6.18 | 99.6 | 17.9 |
| STARCH | %DM | Starch | 2020 | 31.5 | 0.04 | 88.9 | 20.7 |
| SUGARS | %DM | Total sugars | 2123 | 8.0 | 0.05 | 75.6 | 9.9 |

After preliminary exploration, a preprocessing was applied to ***X*** (spectra) consisting to a standard normal variate (SNV) transformation followed by a Savitzky-Golay 2nd derivation (polynomial of order 3, and window of 11 spectral points). Examples of raw and preprocessed spectra are presented in Fig.1. The projected data in PCA scores (Fig.2) illustrates the large spectral heterogeneity existing in the dataset.

**Fig. 1:**
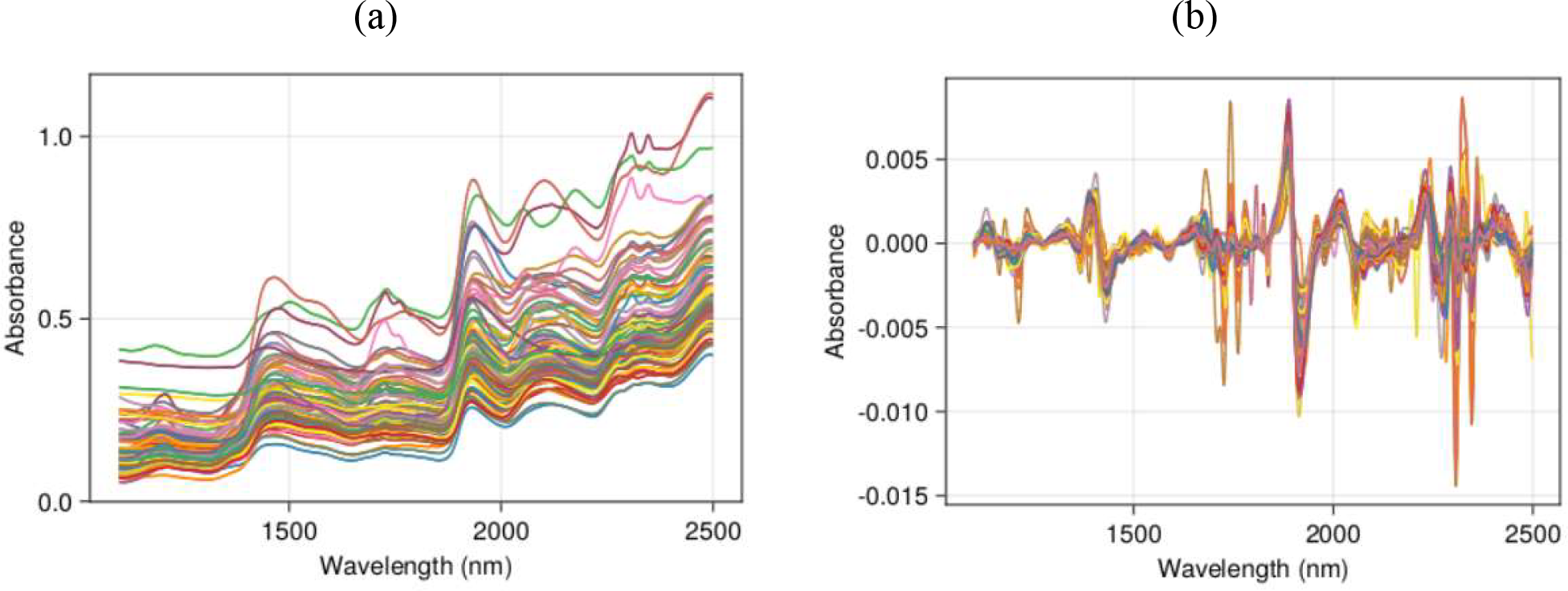
Example of 100 spectra of the dataset: a) raw, b) preprocessed.

### 3.2 Predictive performances of the models

The model performances were compared by computing prediction errors rates on selected test sets (generalization errors). The procedure was the same for each of the compared models and each given response variable (Table 1), as follows. Let us consider the *j*^th^ response variable:

- The total available data for this variable (size *N*_tot,*j*_) is randomly divided to a training set, say *Train* (size *N*_train,*j*_ = 80% * *N*_tot,*j*_), and a test set, say *Test* (size *N*_test,*j*_ = 20% * *N*_tot,*j*_).
- Then, *Train* is randomly divided to a calibration set, say *Cal* (size *N*_cal,*j*_ = 80% *
- *N*_train,*j*_), and a validation set, say *Val* (size *N*_val,*j*_ = 20% * *N*_train,*j*_),
- In summary, *Tot* = *Train* ⏝ *Test*, and *Train* = *Cal* ∪ *Val*.
- The model is tuned by exhaustive grid-search over *Train* as follows:
  - The combinations of the predefined values of the parameters of the model are listed.
  - For each combination of parameter values, the model is fitted on *Cal* and the performance of the combination is evaluated by the root mean squared prediction error rate computed on *Val* (RMSEP_Val_).
- The optimal combination (corresponding to the lower RMSEP_Val_ within the combinations) is then selected and the model is re-fitted (with the selected combination) on *Train* and used to predict *Test*. RMSEP_Test_ is then computed and used as final estimate of generalization error of the given model.

Due to the random splitting (*Train*/*Test* and *Cal*/*Val*), RMSEP_Val_ and RMSEP_Test_ are random variables. Considering only one single replication may lead to possible misleading conclusions since the replication can favor, by chance, one or the other of the models. It is therefore important to replicate the process described in the above items to get the RMSEP distributions. In this article, the process was replicated *n*_rep_ = 30 times. The RMSEP_Test_ distributions were summarized by means and standard errors.

### 3.3 The compared models

The two compared models were the usual kNN-LWPLSR in which the optimal number of LVs, *a*, is determined by CV, and the kNN-LWPLSR-AVG pipeline where a PLSR-averaging is used instead of determining *a*. A particular case of kNN-LWPLSR-AVG was also considered (“omnibus” strategy described thereafter). All the computations were implemented with the package Jchemo [28]. This package is written with the free and fast Julia language [29] (https://julialang.org).

#### kNN-LWPLSR

The preliminary global PLS score space was of dimensionality *nlv*_dis_ = 25 scores, from which were computed, for each new observation to predict, ***x***_new_, the neighborhood (kNN selection based on Mahalanobis distances) and the weights vector *δ*. The grid-search used to tune the model was undertaken over the combinations of the following parameter values, consisting in 315 combinations:

- *h* = {1, 2.5, 5} Shape factor for the weighting function *f*
- *k* = {300, 500, 1000} Neighborhood size (nb. neighbors)
- *nlv* = {0, 1, …, 20} Nb. LVs *a* (LWPLSR on the neighborhood)

#### kNN-LWPLSR-AVG

The same parameter values as above were considered for *h* and *k*, leading to 9 combinations. For each ***x***_new_, the prediction was the average of the predictions of the LWPLSR models (on the neighborhood) with *nlv* = 0, …, 20 LVs (21 predictions).

#### kNN-LWPLSR-AVG with omnibus strategy

When many datasets (spectra × response variables) have to be processed, for instance in automated prediction platforms regularly updated, an “omnibus” strategy applied to PLSR-averaging [19] can facilitate the predictions. This strategy consists in defining a single model (i.e. a set of *a priori* parameter values) that is applied blindly to all the datasets, instead of trying to tune and optimize the model separately for each spectral data and response variable. The defined omnibus model is ideally expected to have an acceptable efficiency for most of the spectral × response variables, even if not always optimal.

In this article, an omnibus strategy was applied to kNN-LWPLSR-AVG and compared to the two above models (kNN-LWPLSR and kNN-LWPLSR-AVG) tuned by grid-search. From preliminary studies on forages (sorghum and other data not considered in this article [8,19]), the omnibus parameters were set to *h* = 1 and *k* = 1000 neighbors.

## 4. Results

The performances of models kNN-LWPLSR and kNN-LWPLSR-AVG tuned by grid-search for the *n*_rep_ = 30 replications are summarized in Fig.3. For all the response variables, the averaging improved the predictions, with the mean RMSEP_Test_ decreasing in relative values from 1.7% (ADF) to 11.4% (SUGARS).

**Fig. 2:**
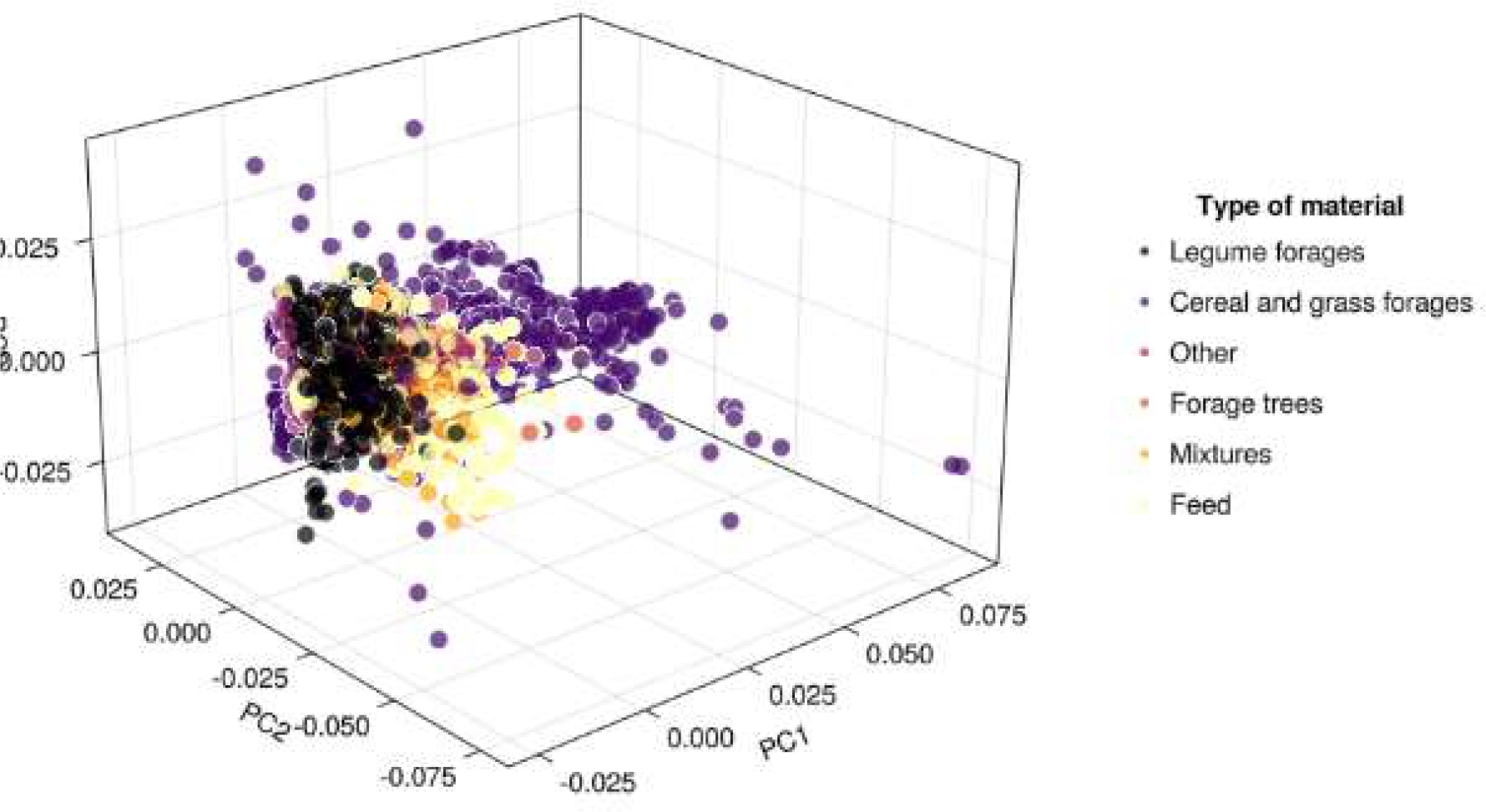
Projection of the observations (preprocessed forage spectra; *N* = 18813) on the three first PCA scores space.

**Figure 3:**
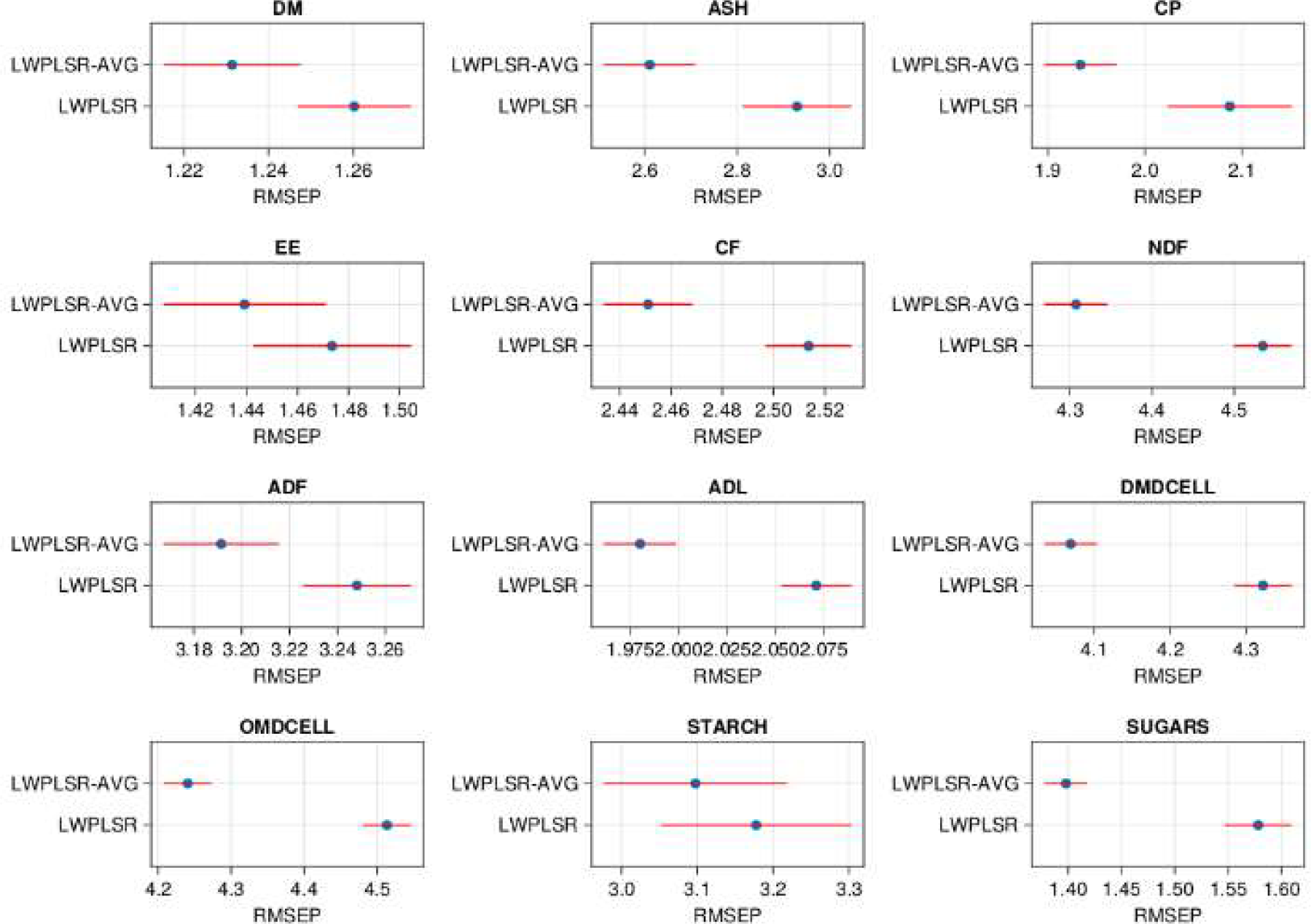
**RMSEP_Test_ for kNN-LWPLSR (“LWPLSR”in the figure) and kNN-LWPLSR-AVG (“LWPLSR_AVG” in the figure) Tuned by grid-search. Averages ± standard errors computed on the *n*rep = 30 replications.**

Fig.4 compares the tuned kNN-LWPLSR-AVG *vs*. the omnibus model (*h* = 1 and *k* = 1000 neighbors). The two strategies showed very close performances: the relative difference in relative value between the mean RMSEP_Test_ were always in the range ± 2.5%. The omnibus model returned lower *vs*. higher error rates depending on the response variables.

**Figure 4:**
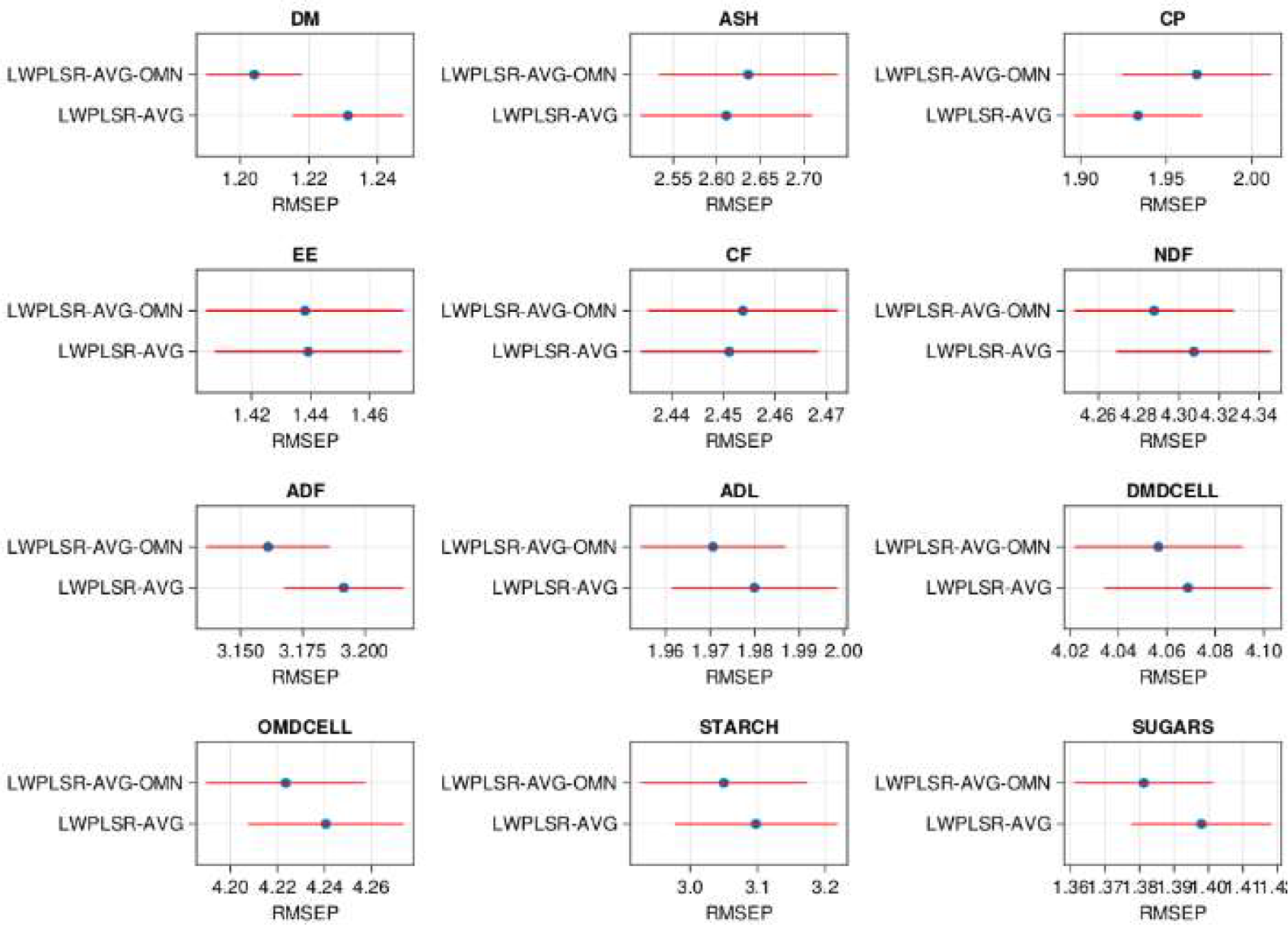
**RMSEP_test_ for kNN-LWPLSR-AVG optimized by grid-search (“LWPLSR_AVG”in the figure) *vs*. with the omnibus model (parameters *h* = 1 and *k* = 100 (“LWPLSR_AVG-OMN” in the figure). Averages ± standard errors computed on the *n*_rep_ = 30 replications.**

## 5. Discussion and conclusions

The averaging always improved the performances of the kNN-LWPLSR models, compared to the usual strategy that optimizes the number of LVs. The improvement was also observed for the omnibus strategy in which single values of parameters *h* and *k* are used for all the response variables. This reinforces the previous results [19] about the interest of using PLS averaging on mixed forage data. Forage and feed spectral data often contain high intrinsic complexity (the material is collected from different species, parts of plants, years, geographical areas, etc.) than can be distributed over all the wavelength range. Averaging different PLS dimensionality may generate better robustness in such situation. The statistical decrease of prediction variances observed with ensemble learning methods [26,30] can also favor better performances.

The present article focused on the uniform weighting, the simplest averaging approach. More elaborated weighting patterns can be considered, but they seem not be always more efficient [19]. Moreover, in the context of local PLSR, they can become highly time consuming to implement on data. For instance, additional computations on the study dataset (not detailed in this article) showed that a weighting based on the CV-errors [19] did not improve the predictions. And a weighting based on AIC [19] required several days of computation to run the *n*_rep_ = 30 replications, which is not realistic in practice. The uniform weighting seems therefore to be a good compromise for kNN-LWPLSR-AVG.

To reinforce the finding of this article, kNN-LWPLSR-AVG should be evaluated on other agronomic materials than mixed forages and feed, with the same or, conversely, lower level of heterogeneity (data complexity). In the second case, the method may be less advantageous. In practice, such evaluations can easily be implemented either by using already existing routines (e.g. package Jchemo [28]) or by inserting the averaging in other available pipelines of local PLSR.

## Acknowledgements

I thank Laurent Bonnal (Cirad, Montpellier) for the preparation of the spectral and chemical data.

## Notes

https://github.com/mlesnoff/Jchemo.jl

### Competing Interest Statement

The authors have declared no competing interest.

### Summary of Updates

The manuscript text was improved.

## References

[1] A. Höskuldsson, PLS regression methods, Journal of Chemometrics. 2 (1988) 211–228. 10.1002/cem.1180020306.

[2] H. Wold, Nonlinear iterative partial least squares (NIPALS) modeling: some current developments, in: Multivariate Analysis II, Krishnaiah, P. R., Wright State University, Dayton, Ohio, USA. June 19–24, 1972. New York: Academic Press, 1973: pp. 383–407.

[3] S. Wold, M. Sjöström, L. Eriksson, PLS-regression: a basic tool of chemometrics, Chemometrics and Intelligent Laboratory Systems. 58 (2001) 109–130. 10.1016/S0169-7439(01)00155-1.

[4] P. Dardenne, G. Sinnaeve, V. Baeten, Multivariate Calibration and Chemometrics for near Infrared Spectroscopy: Which Method?, J. Near Infrared Spectrosc., JNIRS. 8 (2000) 229–237.

[5] M. Clairotte, C. Grinand, E. Kouakoua, A. Thébault, N.P.A. Saby, M. Bernoux, B.G. Barthès, National calibration of soil organic carbon concentration using diffuse infrared reflectance spectroscopy, Geoderma. 276 (2016) 41–52. 10.1016/j.geoderma.2016.04.021.

[6] F. Davrieux, D. Dufour, P. Dardenne, J. Belalcazar, M. Pizarro, J. Luna, L. Londoño, A. Jaramillo, T. Sanchez, N. Morante, F. Calle, L. Becerra Lopez-Lavalle, H. Ceballos, LOCAL regression algorithm improves near infrared spectroscopy predictions when the target constituent evolves in breeding populations, Journal of Near Infrared Spectroscopy. 24 (2016) 109. 10.1255/jnirs.1213.

[7] H. Tran, P. Salgado, E. Tillard, P. Dardenne, X.T. Nguyen, P. Lecomte, “Global” and “local” predictions of dairy diet nutritional quality using near infrared reflectance spectroscopy, Journal of Dairy Science. 93 (2010) 4961–4975. 10.3168/jds.2008-1893.

[8] M. Lesnoff, M. Metz, J.-M. Roger, Comparison of locally weighted PLS strategies for regression and discrimination on agronomic NIR data, Journal of Chemometrics. n/a (2020) e3209. 10.1002/cem.3209.

[9] J.A. Fernández Pierna, A. Laborde, L. Lakhal, M. Lesnoff, M. Martin, Y. Roggo, P. Dardenne, The applicability of vibrational spectroscopy and multivariate analysis for the characterization of animal feed where the reference values do not follow a normal distribution: A new chemometric challenge posed at the ‘Chimiométrie 2019’ congress, Chemometrics and Intelligent Laboratory Systems. 202 (2020) 104026. 10.1016/j.chemolab.2020.104026.

[10] A.A. Gowen, G. Downey, C. Esquerre, C.P. O’Donnell, Preventing over-fitting in PLS calibration models of near-infrared (NIR) spectroscopy data using regression coefficients, Journal of Chemometrics. 25 (2011) 375–381. 10.1002/cem.1349.

[11] J.H. Kalivas, Multivariate Calibration, an Overview, Analytical Letters. 38 (2005) 2259–2279. 10.1080/00032710500315904.

[12] H. van der Voet, Comparing the predictive accuracy of models using a simple randomization test, Chemometrics and Intelligent Laboratory Systems. 25 (1994) 313–323. 10.1016/0169-7439(94)85050-X.

[13] F. Westad, F. Marini, Validation of chemometric models – A tutorial, Analytica Chimica Acta. 893 (2015) 14–24. 10.1016/j.aca.2015.06.056.

[14] S. Wiklund, D. Nilsson, L. Eriksson, M. Sjöström, S. Wold, K. Faber, A randomization test for PLS component selection, Journal of Chemometrics. 21 (2007) 427–439. 10.1002/cem.1086.

[15] M. Lesnoff, J.-M. Roger, D.N. Rutledge, Monte Carlo methods for estimating Mallows’s Cp and AIC criteria for PLSR models. Illustration on agronomic spectroscopic NIR data, Journal of Chemometrics. n/a (2021) e3369. 10.1002/cem.3369.

[16] D.D. Silalahi, H. Midi, J. Arasan, M.S. Mustafa, J.-P. Caliman, Automated Fitting Process Using Robust Reliable Weighted Average on Near Infrared Spectral Data Analysis, Symmetry. 12 (2020) 2099. 10.3390/sym12122099.

[17] M.H. Zhang, Q.S. Xu, D.L. Massart, Averaged and weighted average partial least squares, Analytica Chimica Acta. 504 (2004) 279–289. 10.1016/j.aca.2003.10.056.

[18] J. Shenk, M. Westerhaus, P. Berzaghi, Investigation of a LOCAL calibration procedure for near infrared instruments, Journal of Near Infrared Spectroscopy. 5 (1997) 223. 10.1255/jnirs.115.

[19] M. Lesnoff, D. Andueza, C. Barotin, P. Barre, L. Bonnal, J.A. Fernández Pierna, F. Picard, P. Vermeulen, J.-M. Roger, Averaging and Stacking Partial Least Squares Regression Models to Predict the Chemical Compositions and the Nutritive Values of Forages from Spectral Near Infrared Data, Applied Sciences. 12 (2022) 7850. 10.3390/app12157850.

[20] S. Schaal, C.G. Atkeson, S. Vijayakumar, Scalable Techniques from Nonparametric Statistics for Real Time Robot Learning, Applied Intelligence. 17 (2002) 49–60. 10.1023/A:1015727715131.

[21] S. Kim, M. Kano, H. Nakagawa, S. Hasebe, Estimation of active pharmaceutical ingredients content using locally weighted partial least squares and statistical wavelength selection, International Journal of Pharmaceutics. 421 (2011) 269–274. 10.1016/j.ijpharm.2011.10.007.

[22] E. Sicard, R. Sabatier, Theoretical framework for local PLS1 regression, and application to a rainfall data set, Computational Statistics & Data Analysis. 51 (2006) 1393–1410. 10.1016/j.csda.2006.05.002.

[23] W.S. Cleveland, Robust Locally Weighted Regression and Smoothing Scatterplots, Journal of the American Statistical Association. 74 (1979) 829. 10.2307/2286407.

[24] W.S. Cleveland, S.J. Devlin, Locally Weighted Regression: An Approach to Regression Analysis by Local Fitting, Journal of the American Statistical Association. 83 (1988) 596–610. 10.1080/01621459.1988.10478639.

[25] B.S. Dayal, J.F. MacGregor, Improved PLS algorithms, Journal of Chemometrics. 11 (1997) 73–85. 10.1002/(SICI)1099-128X(199701)11:1<73::AID-CEM435>3.0.CO;2-#.

[26] T. Hastie, R. Tibshirani, J. Friedman, The elements of statistical learning: data mining, inference, and prediction, 2nd ed., Springer, New York, 2009.

[27] K.P. Burnham, D.R. Anderson, Model selection and multimodel inference: a practical information-theoretic approach, 2nd ed., Springer, New York, NY, USA, 2002.

[28] M. Lesnoff, Jchemo: Julia package for machine learning and chemometrics on highdimensional data, (2021). https://github.com/mlesnoff/Jchemo.

[29] J. Bezanson, A. Edelman, S. Karpinski, V.B. Shah, Julia: A fresh approach to numerical computing, SIAM Rev. 59 (2017) 65–98. 10.1137/141000671.

[30] K.P. Burnham, D.R. Anderson, Multimodel Inference: Understanding AIC and BIC in Model Selection, Sociological Methods & Research. 33 (2004) 261–304. 10.1177/0049124104268644.

